# Plasma membrane H^+^-ATPase activation increases global transcript levels and promotes the shoot growth of light-grown Arabidopsis seedlings

**DOI:** 10.1101/2024.08.30.610460

**Authors:** Satoru Naganawa Kinoshita, Kyomi Taki, Fumika Okamoto, Mika Nomoto, Koji Takahashi, Yuki Hayashi, Junko Ohkanda, Yasuomi Tada, Iris Finkemeier, Toshinori Kinoshita

**Affiliations:** Institute of Plant Biology and Biotechnology, University of Münster, Germany; Graduate School of Science, Nagoya University, Japan; Institute of Agriculture Shinshu University, Japan; Center for Gene Research, Nagoya University, Japan; ITbM, Nagoya University, Japan

**Keywords:** Acid growth, Arabidopsis thaliana, Ca^2+^ signalling, PM H^+^-ATPase, shoot growth, transcriptomics

## Abstract

(1) Plant cell growth require the elongation of cells mediated by cell wall remodelling and turgor pressure changes. The plasma membrane (PM) H^+^-ATPase facilitates both cell wall remodelling and turgor pressure changes, by acidifying the apoplast of cells, referred to as acid growth. The acid growth theory is mostly established on the auxin-induced activation of PM H^+^-ATPase in non-photosynthetic tissues. However, how PM H^+^-ATPase affect the growth in photosynthetic tissues of Arabidopsis remains unclear.
(2) Here, a combination of transcriptomics and cis-regulatory element analysis was conducted to identify the impact of PM H^+^-ATPase on global transcript levels and the molecular mechanism downstream of the PM H^+^-ATPase.
(3) The PM H^+^-ATPase activation increased transcript levels globally, especially cell wall modification-related genes. The transcript level changes were in PM H^+^-ATPase-dependent manner. Involvement of Ca^2+^ was suggested as CAMTA motif was enriched in the promoter of PM H^+^-ATPase-induced genes and cytosolic Ca^2+^ elevated upon PM H^+^-ATPase activation.
(4) PM H^+^-ATPase activation in photosynthetic tissues promote the expression of cell wall modification enzymes and shoot growth, adding a novel perspective of photosynthesis-dependent PM H^+^-ATPase activation in photosynthetic tissues to the acid growth theory that has primarily based on findings from non-photosynthetic tissues.

## Introduction

Plant growth is mediated by the expansion and division of individual cells. Expansion and division processes are linked to the structure and property of the cell wall, a rigid yet flexible barrier surrounding plant cells. The dynamic equilibrium between rigidity and flexibility is crucial for plant development (Cooper, 2000). The remodelling of the cell wall is mediated by a decrease in apoplastic pH, activating a cascade of enzymatic reactions that loosen the cell wall structure. The apoplastic acidification model is also called acid growth model. The acid growth model has been based on extensive studies from 1970s (Rayle & Cleland, 1970; Hager *et al*., 1971) and the model illustrates that the phytohormone auxin induces the apoplastic acidification by activating plasma membrane (PM) P-type H^+^-ATPase, a primary source of H^+^ gradient across plasma membrane (Palmgren, 2001; Hager, 2003; Takahashi *et al*., 2012). An application of an unique fungal phytotoxin Fusicoccin, an irreversible activator of PM H^+^-ATPase, has revealed that sole activation of PM H^+^-ATPase can achieve the growth of cells, further illustrating the central role of PM H^+^-ATPase in the acid growth model (Kutschera & Schopfer, 1985). Other hormones also control the apoplast acidification and cell growth, i.e. brassionsteroids (BRs) promote PM H^+^-ATPase activity (Minami *et al*., 2019).

Phytohormone-mediated PM H^+^-ATPase activity regulation consists of two distinct mechanisms. Upon the perception of the phytohormone, both types of regulation are achieved by posttranslational modification of PM H^+^-ATPase C-terminal auto-inhibitory domain (Miao *et al*., 2022). The C-terminal auto-inhibitory domain have several phosphorylation sites that control the activity of the PM H^+^-ATPase (Falhof *et al*., 2016; Fuji *et al*., 2024; Hayashi *et al*., 2024). Taking auxin as one example of regulation of PM H^+^-ATPase C-terminal penultimate Thr phosphorylation, perception of auxin at the PM is mediated by auxin binding protein (ABP) and PM-localised transmembrane kinases (TMKs), phosphorylating PM H^+^-ATPase (Li *et al*., 2021b; Lin *et al*., 2021; Friml *et al*., 2022). In addition, the perception of the auxin in the nucleus is mediated by TIR1/AFB complex, which induces the expression of *small auxin up RNAs* (*SAURs*) family genes (Ang & Østergaard, 2023). SAURs proteins interact with protein phosphatase 2C D-clade (PP2C-D) family proteins and inhibit the phosphatase activity of PP2C-D, maintaining the phosphorylation status of PM H^+^-ATPase (Ren & Gray, 2015).

Other than phytohormonal regulation, recent studies in photosynthetic tissues have discovered that photosynthesis promotes the activation of the PM H^+^-ATPase via phosphorylation of C-terminus residues (Okumura *et al*., 2016; Kinoshita *et al*., 2023; Hayashi *et al*., 2024). The activation of the PM H^+^-ATPase in leaves is dependent on the photosynthetic activity and the photosynthesis product, but independent from the light signalling mediated by known photoreceptor, i.e., phytochrome, phototropin, nor cryptochrome (Okumura *et al*., 2016). Further investigation on activation mechanism of photosynthesis-dependent PM H^+^-ATPase has found a novel activator of PM H^+^-ATPase in light illuminated and sugar fed leaves, SAUR30 (Kinoshita *et al*., 2023), that is not responsive to external auxin application (Paponov *et al*., 2008). The regulatory mechanism of PM H^+^-ATPase by photosynthesis emerges a fundamental question: What physiological roles does photosynthesis-dependent PM H^+^-ATPase activation play in photosynthetic tissues? Nitrate uptake into leaves has been proposed as one role of the photosynthesis-dependent activation of PM H^+^-ATPase (Kinoshita *et al*., 2023). In addition, it is noteworthy that an acid growth model has been established in divergent plant species, primarily using coleoptile, hypocotyl of etiolated seedlings or roots, non-photosynthetic tissues. However, the growth of cells in photosynthetic tissues such as leaves and light-grown seedlings, green seedlings, has a limited investigation; in other words, how PM H^+^-ATPase activity promotes growth in photosynthetic tissues remains elusive. In addition, while transcriptional inhibition is suggested to supresses the acid growth of cells (Arsuffi & Braybrook, 2018), the impact of PM H^+^-ATPase activation alone—excluding light or phytohormone—on global transcript levels in photosynthetic tissues remains unknown.

Here, we investigated the impact of PM H^+^-ATPase activation on the growth of green seedlings and the global transcriptome, finding that PM H^+^-ATPase activation promotes the growth of green seedlings via upregulation of cell wall-related gene expression. Further *in silico* analysis using the promoters of PM H^+^-ATPase activation-responsive genes predicted the involvement of Ca^2+^ signalling, that was confirmed with observation using the Ca^2+^ biosensor, GCaMP3 (Toyota *et al*., 2018). From the results, we propose a novel perspective in the acid growth model, indicating that PM H^+^-ATPase activation in photosynthetic tissues promote the growth of cells by directly or indirectly inducing Ca^2+^ signalling and remodelling the cell wall properties. Considering the PM H^+^-ATPase is activated mainly by photosynthesis and photosynthetic products, our findings in photosynthesis tissues suggest one of the physiological roles of photosynthesis-dependent activation of PM H^+^-ATPase, shedding light on a novel viewpoint of photosynthesis in the framework of the acid growth theory.

## Materials and Methods

### Plant materials and growth

The dicot model plant *Arabidopsis thaliana* accession Columbia-0 (Col-0) was used as wildtype, except for the hypocotyl length measurement of *open stomata2-1D (ost2-1D)* using Landsberg *erecta* (Ler) as background wildtype. Seeds were sterilised and stratified with Plant preservative mixture (PPM)-tween solution [2% (v/v) PPM (Plant Cell Technology), 0.005% (v/v) Tween-20 (Fujifilm)] at 4°C for 2 nights. After washing, the seeds were sown on 0.8% agar Murashige and Skoog (MS) plate [1/2 MS salt, 0.8% (w/v) Agar (Fujifilm), 2.3 mM MES-KOH pH 5.7 (Nacalai)] or 0.6% Gellan gum MS [1/2 MS salt, 0.6% (w/v) Gellan Gum (Fujifilm), 2.3 mM MES-KOH pH 5.7 (Nacalai)] for RNA-seq samples or other experiments, respectively. The plants were grown under long day cycle, 16h light: 8h dark (6:00 h to 22:00 h light at Japan standard time, JST) at 22−23°C with a photon flux 40−70 µmol m^−2^ s^−1^ of white light. The plates were put horizontally on the shelf to prevent the seedling hypocotyl from touching the plate. The 7-day-old seedlings were collected for hypocotyl measurement or transferred to darkness before the treatment of Fusicoccin or EtOH.

### Hypocotyl length measurements

The 7-day-old seedlings of Col-0, *aha1-9* (SAIL_1285_D12; Yamauchi *et al*., 2016), complementation line *gAHA1*/ *aha1-9* (Hayashi *et al*., 2024), Ler, and *ost2-1D*/ Ler (Merlot *et al*., 2002, 2007) were transferred and lined onto the agar plate, followed by photograph. The hypocotyl lengths of each genotype in images were manually measured by Fiji software (ImageJ v.1.54).

### Extraction of Fusicoccin-A

Fusicoccin-A (Fc-A) was extracted from *Phomopsis amygdali Niigata* 2(Sassa *et al*., 1999), following the procedure in (Sassa *et al*., 2002). Products were extracted with ethyl acetate, and the crude materials were separated by silica gel column chromatography. Further purification by recrystallization from ethyl acetate gave FC-A as a colourless powder, purity > 95% on NMR (Ohkanda *et al*., 2023).

### Treatment on seedling shoot

Plates with 7-day-old seedlings were put in a box and kept in the dark for overnight before the treatments to reduce the PM H^+^-ATPase activity. As one biological replicate, 20 seedling shoots were separated from the roots and incubated in the 12-well plate filled with 1/2 MS liquid media [1/2 MS salt, 2.3 mM MES-KOH, pH 5.7 except for low pH treatment] containing treatments at final concentration of 0.1% (v/v) ethanol (EtOH), 30 µM Fc-A in 0.1% ethanol, or 1/2 MS liquid media with 2 mM MES-KOH pH 4.0 (low pH treatment) for 2 hours in the dark. The samples were then flash frozen in liquid nitrogen and kept in −80°C until the RNA extractions.

### RNA extraction and cDNA synthesis

Frozen samples were homogenised by beads, and total RNA was purified using the NucleoSpin RNA Plant (MACHEREY-NAGEL) kit, following the manufacturer’s instructions. For real time-quantitative PCR, complementary DNAs (cDNA) were synthesised from 400 ng of purified total RNAs using the PrimeScriptRT Reagent Kit (TaKaRa).

### Preparation of complementary DNA libraries and RNA-seq analysis

For preparation of cDNA libraries, the quality of total RNAs were analysed by the Agilent 4150 TapeStation System (Agilent) and then mRNAs were enriched from 600 ng of total RNAs, using Next poly(A) mRNA Magnetic Isolation Module (New England Biolabs, NEB), followed by first-strand cDNA synthesis with the Next Ultra II RNA Library Prep Kit for Illumina (NEB) and Next Multiplex Oligo for Illumina (NEB). cDNA libraries were then sequenced as single end reads for 81 bp on NextSeq 550 system (Illumina). For each treatment condition, three independent biological replicates were sequenced individually and 4.2−6.3 million total reads were obtained. The sequence reads were then mapped to Arabidopsis thaliana reference genome, TAIR10 (Ensemble), with the function “RNA-seq Alignment” in the web platform, Illumina Basespace (Illumina). Normalisation and statistical analysis for differentially expressed genes were conducted in the web platform, Degust v3.2.0 (Powell, 2019). The differentially expressed genes were defined as log_2_(|foldchange|)> 1 and false discovery rate (FDR) =< 0.05, compared to EtOH control treatment. For the visualisation of functionally grouped gene expression, MapMan ontology (Thimm *et al*., 2004) was imported from “Ath_AGI_LOCUS_TAIR10_Aug2012.txt” on the website.

### Real time-quantitative PCR

Primers for amplifying the target genes and internal standard gene were listed in **Table S1**. Primers and cDNA were mixed with SYBR Green PCR Master Mix (Applied Biosystems) and the fluorescent signals along the cycles were quantified in StepOnePlus™ Real-time PCR System (Applied Biosystems). The data obtained from the systems was further analysed in Real-time PCR Miner web platform (Zhao & Fernald, 2005) to calculate the corrected Ct values. The Ct values of target genes were normalised by the Ct value of internal standard gene, *UBQ5*, to calculate the relative expression of each gene.

### *In silico* promoter analysis

The statistical analysis for overrepresented pentamers in the promoter of the top 200 most Fc-A- and low pH-induced genes were conducted by following the pentamer prediction program developed in (Yamamoto *et al*., 2011). The top 4 most overrepresented pentamers with upstream and downstream were extracted from the datasets (top 200 Fc-A- and low pH-induced genes) to visualise the frequency of the octamer sequence on the WebLogo web platform (https://weblogo.berkeley.edu/logo.cgi).

### Cytosolic Ca^2+^ imaging

The plate with 7-day-old seedlings of Ca^2+^ biosensor line, p35S:GCaMP3 plant (Toyota *et al*., 2018; Matsumura *et al*., 2022), were put in a box and kept in the dark overnight. In a dark room, seedlings were transferred to the stage of motorised fluorescence stereomicroscope (SMZ-25; Nikon) with a 1× objective lens (NA= 0.156, P2-SHR PLAN APO; Nikon). Live imaging of GCaMP3 fluorescence was conducted using the sCMOS camera (ORCA-Fusion BT; Hamamatsu Photonics) attached to the stereomicroscope with the NIS-Elements software (Nikon). After adjusting the focus on the seedling hypocotyl, liquid ½ MS media with 0.1% EtOH or 30 μM Fc-A was gently applied to the neighbouring region of the seedling, avoiding mechanical contact with the seedling. The fluorescence intensities were quantified using Fiji software (ImageJ v.1.54). The fractional fluorescence changes (ΔF/F_0_) were calculated using ΔF/F_0_ = (F−F_0_) /F_0_, where F_0_ is the average baseline fluorescence determined by the average of fluorescence intensities in the first 5 min.

### Data analysis, statistics, and graph generation

Most data transformation, statistical analysis, and graph generation were performed in Rstudio (R version 4.3.1) with the packages Tidyverse, multicomp, and ggplot2, except for the MapMan ontology. The *P* value for the MapMan ontology was calculated using the Whitney U test in the MapMan software.

### Accession numbers

AHA1, AT2G18960; XTH23, AT4G25810; AGP18, AT4G37450; EXPL2, AT4G38400; PAE11, AT5G45280; TCH4, AT5G57560; SAUR30, AT5G53590

## Results

### PM H^+^-ATPase promoted the growth of green seedlings

To test whether PM H^+^-ATPase activity influences the shoot growth of green seedlings, as an indicator of the seedling growth, the hypocotyl length in loss-of-function mutant and constitutive active mutant of one of two major PM H^+^-ATPase, *Arabidopsis H^+^-ATPase1* (*AHA1*), was measured. The loss-of-function mutant, *aha1-9*, showed reduced growth compared to wildtype and the growth was recovered by the complementation of *genomic AHA1* in *aha1-9* (**Fig 1A**, **Fig S1**). On the contrary, the AHA1 constitutive active mutant, *open stomata 2-1 dominant* (*ost2-1D*), showed a longer hypocotyl length compared to wildtype (**Fig 1B**). These results illustrated that the PM H^+^-ATPase activity plays a significant role in the growth of green seedlings.

**Figure 1.**
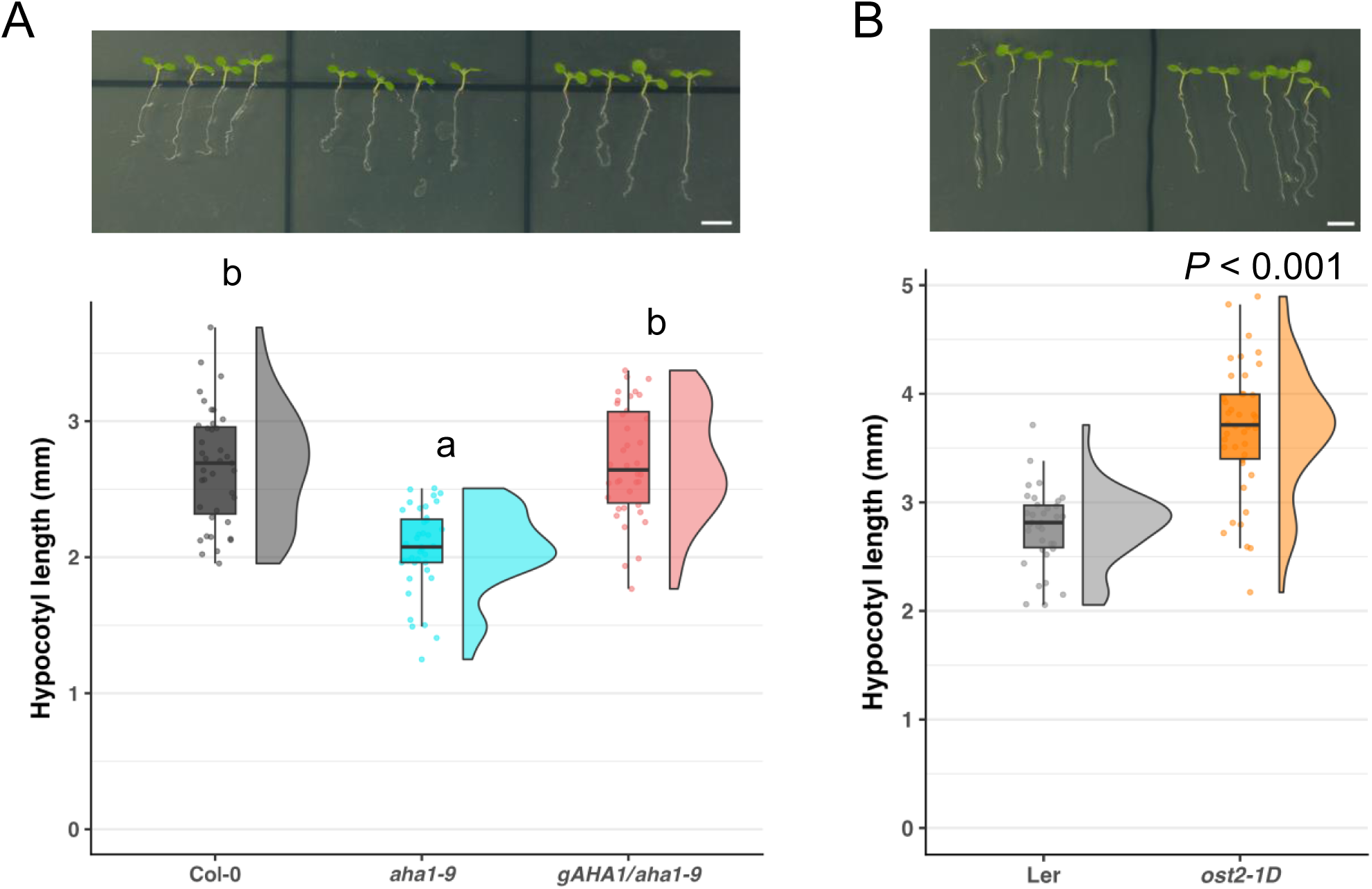
Hypocotyl elongation of seedlings with different PM H^+^-ATPase activities. A) Hypocotyl length of one-week-old wildtype (Columbia-0, Col-0), *aha1-9*, complementation line *gAHA1*/*aha1-9.* Each dot indicate the length of one seedling, and the boxplot and half-violin plot present the distribution and density of hypocotyl length across the genotypes; *n* = 38−39. Different letters above boxplot indicate the significant difference, determined by one-way ANOVA with Tukey HSD. B) Hypocotyl length of one-week-old wildtype (Landsberg electra, Ler) and *aha1* mutant with constitutive activation mutation (*open stomata 2-1 dominant*, *ost2-1D).* Each dot indicates the length of one seedling and the boxplot and half-violin plot represent the distribution and density of hypocotyl length across the genotypes; *n* = 34−42. The *P* value was determined by Welch’s *t* test. Scale bars: 5 mm

### Global transcript expression was affected by PM H^+^-ATPase activation

To investigate the mechanism of PM H^+^-ATPase-dependent growth in green seedlings, the changes in transcript levels were detected by RNA-seq. Seedlings were kept in the dark overnight to reduce the activity of PM H^+^-ATPase (Okumura *et al*., 2016; Kinoshita *et al*., 2023) and then incubated in liquid media with ethanol (EtOH) or a PM H^+^-ATPase activator, Fusicoccin-A (Fc-A) (**Fig 2A**). Fc-A causes the activation of PM H^+^-ATPase via glueing the interaction of 14-3-3 protein and penultimate threonine residue (Thr) of PM H^+^-ATPase C-term (Fuglsang *et al*., 2003; Ohkanda *et al*., 2023). Due to the high price of commercially available Fc-A that is unsuitable for mass consumption, this study extracted and purified Fc-A from a fungus, *Phomopsis amygdali* (Sassa *et al*., 2002). Comparing ethanol- and Fc-A-treated seedlings in the dark can elucidate the transcriptional changes downstream of PM H^+^-ATPase activation in the seedlings. Since the activation of PM H^+^-ATPase lowers pH in the apoplast, seedlings treated by low pH liquid media were included for the RNA-seq analysis. Interestingly, activation of PM H^+^-ATPase by Fc-A induced considerable changes in transcript levels compared to the low pH treatments (**Fig 2B** and **Data S1**), implying the PM H^+^-ATPase activation stimulated the divergent response in plant cells. Transcript levels in some gene functional groups determined by MapMan were significantly different, especially the genes that have roles in signalling, cell wall, and stress (**Fig 2C**). A more specific grouping of the genes showed the genes involved in cell wall modification (**Fig S2**). Since Fc-A is a fungal toxin, exogenous application of Fc-A may induce the artefact effect on the transcriptional changes. To investigate the impact of PM H^+^-ATPase activation in green tissues from other perspective, previous transcriptome analyses from light-illuminated and sucrose-fed leaves were compared as a reference because light illumination and sucrose feeding also induce PM H^+^-ATPase activity (Kinoshita *et al*., 2023). GO enrichment analysis on the transcripts that were induced by Fc-A treatment in seedlings, light illumination and sucrose feeding on leaves showed the significant enrichment in genes involved in cell wall biogenesis and glucosyl modification (**Fig S3**), suggesting that the PM H^+^-ATPase activation in green tissues promotes the transcript level of cell wall-related genes and therefore the promote the cell growth by modifying the cell wall property. It should be noted that similar GO enrichment in cell wall modification genes has been observed in the low pH treated root of Arabidopsis (Lager *et al*., 2010). However, our transcriptome analysis identified specific genes in green seedlings or novel genes involved in PM H^+^-ATPase activation-dependent cell wall genes (**Fig S4**).

**Figure 2.**
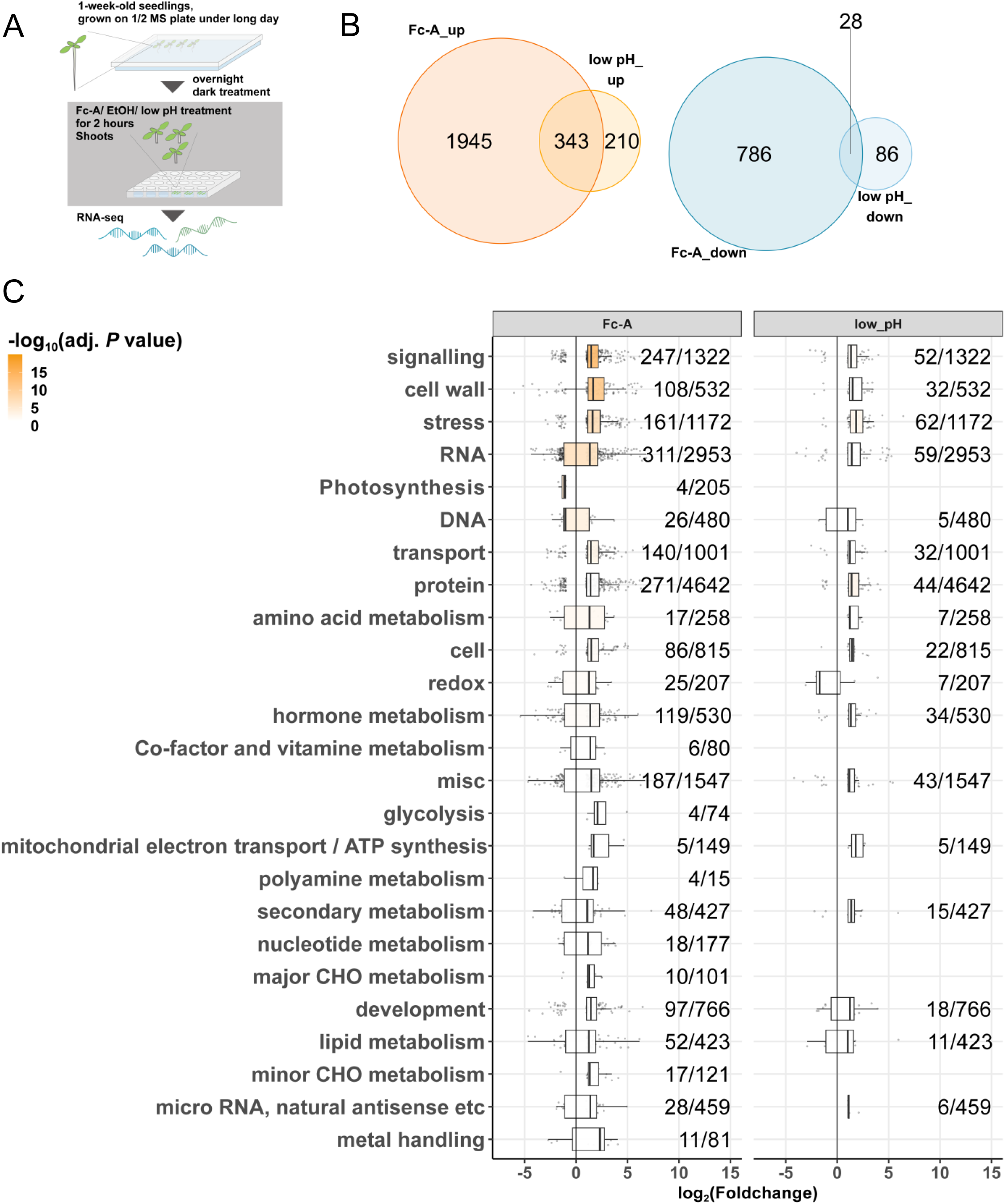
Transcriptome changes in Fc-A-treated seedling shoots. A) Schematic procedure of treatment and sampling conditions. One-week-old seedlings were transferred to the darkness for overnight and then the above-ground part of seedlings were incubated in ½ MS liquid media with 30 µM Fc-A or equal amount of ethanol for 2 hours. In addition, as the low pH treatment, the seedling shoots were kept in pH4.0 buffered ½ MS liquid medium for 2 hours. B) Venn diagrams represent the numbers of differentially expressed genes (DEGs; foldchange > 2, FDR < 0.05) in Fc-A treated or low pH media treated seedlings compared to ethanol treated seedlings. On the left are upregulated DEGs and on the right are downregulated DEGs. C) The boxplots represent the distribution of DEGs expression changes in functional category groups compared to the ethanol treated samples. The numbers beside the boxplots indicate the number of DEGs over the number of all registered genes in the category groups. Adjusted *P* values were determined by Whitney U test with in MapMan.

### Cell wall modification-related genes were induced in PM H^+^-ATPase activity-dependent manner

To further confirm whether the cell wall modification genes induced by Fc-A depend on PM H^+^-ATPase, the changes in transcript levels were checked by RT-qPCR. As expected, the induction of all tested cell wall-related genes was significantly reduced in the *aha1-9* mutant upon Fc-A treatment (**Fig 3**), indicating that the AHA1 plays an essential role in inducing the cell wall-related gene expression. In addition, Fc-A transcriptome analysis revealed that the expression level of PM H^+^-ATPase activator, *Small Auxin Up RNA 30* (*SAUR30*), was induced by Fc-A treatment, and the *SAUR30* gene expression induction by Fc-A was reduced in *aha1-9* (**Fig S5A**). A recent study has revealed that SAUR30 is a positive regulator in photosynthetic product-dependent activation of PM H^+^-ATPase (Kinoshita *et al*., 2023). Our results suggest that SAUR30 regulates PM H^+^-ATPase in a positive feedback manner.

**Figure 3.**
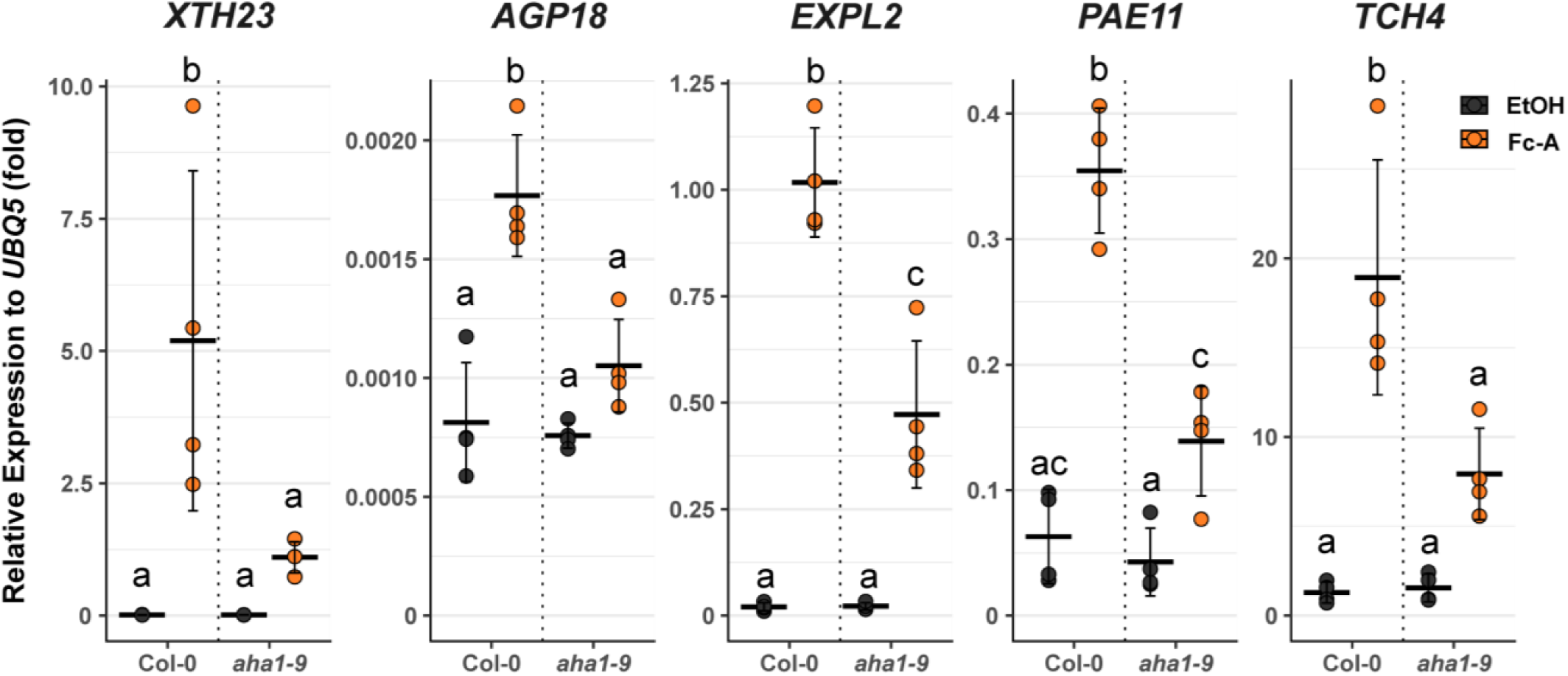
PM H+-ATPase-dependent expression changes of cell wall-related genes. The relative expression of representative cell wall-related genes to ubiquitous *UBQ5* in wildtype (Col-0) and *aha1-9,* determined by RT-qPCR. Each point represents one biological replicates. Crossbars and error bars represent the mean ± S.D. in each conditions. Different letters above error bars indicate the significant difference in each gene expression, determined by one-way ANOVA with Tukey HSD.

### Putative CAMTA binding pentamer was enriched in the promoters of PM H^+^-ATPase- and low pH-driven DEGs

To investigate how PM H^+^-ATPase activation affects the global transcript levels, 1000 bp upstream of transcription start site of the genes that were highly increased upon Fc-A and low pH medium treatment were extracted from the database and then computationally analysed how frequently specific pentamer can be found in the datasets (Matsumura *et al*., 2022). The input genes were the top 200 most Fc-A-induced genes out of genes that were increased upon Fc-A and low pH treatment. Interestingly, the three most frequently observed pentamers in the tested promoters were similar to the Ca^2+^ transcription factor CAMTA binding cis-elements, CACGC or CGCGT (complementary, ACGCG) (**Fig 4A**). Most tested cell wall-related genes had the same pentamers in their 1000 bp upstream of 5’-UTR, except TCH4 had the pentamer within 5’-UTR (**Fig 4B**) as well as SAUR30 (**Fig S5B**). In addition, the comparison of our transcriptomics results with the previously detected cell wall-related gene expression in *camta* mutants over wildtype showed mostly reversed patterns (**Fig S6;** Kim *et al*., 2013), and Fc-A upregulated transcripts overlapped CAMTA regulated touch-induced transcripts (**Fig S7**; Darwish *et al*., 2022). Those results suggest that the Fc-A treatment involves the activation of CAMTA and, therefore, induces the expression of various genes including cell wall-related genes, implying that the Ca^2+^ signalling and CAMTA transcription factors may be involved in the global transcript level changes by PM H^+^-ATPase activation.

**Figure 4.**
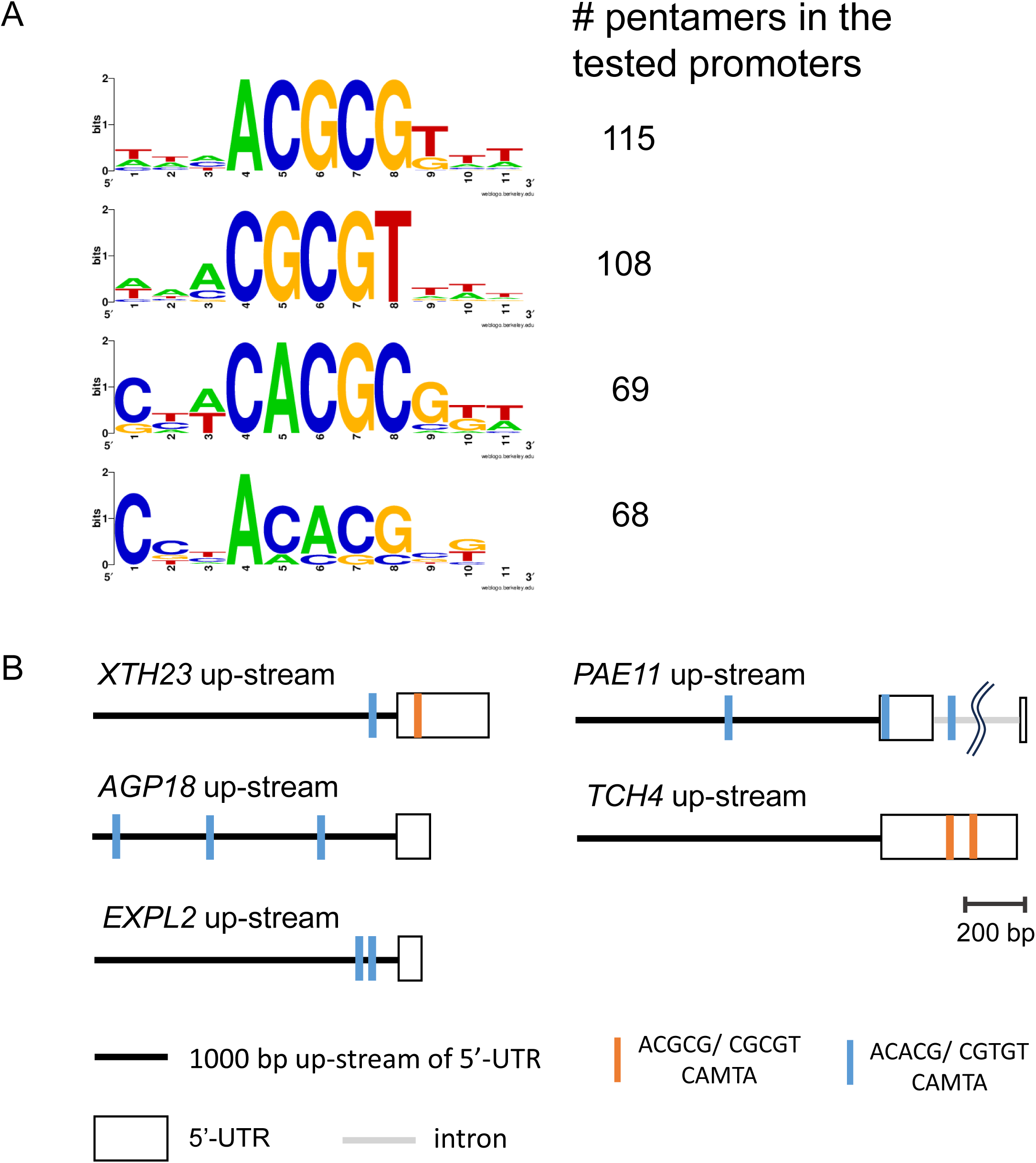
Frequently appeared pentamer in Fc-A and low pH treatment. A) The frequently observed pentamers were searched in the 1000 bp upstream of TOP200 most Fc-A & low pH-induced genes’ 5’-UTRs. Numbers next to the motif logos represent the numbers of pentamers in the datasets. Top 4 most observed pentamers, mostly CAMTA-binding motif GCGC box [CGC(/T)GT], are listed. B) Schematic diagram of the pentamer positions in the promoter region of cell wall-related genes. Black line represent the 1000 bp upstream of 5’-UTR; blue narrow line and orange narrow line indicate the position of GCGC box, ACGCG and ACACG pentamer, respectively; white box represent the position of 5’-UTR.

### PM H^+^-ATPase activation increased cytosolic Ca^2+^ levels in the hypocotyl of green seedlings

To test whether PM H^+^-ATPase activation induces the Ca^2+^ elevation in the cytosol as the CAMTA activation requires cytosolic Ca^2+^ elevation, the plants expressing a well-established Ca^2+^ biosensor, GCaMP3 were treated with Fc-A or ethanol, and the GCaMP3 fluorescence was monitored. Monitoring the GCaMP3 biosensor revealed that the cytosolic Ca^2+^ levels gradually increased upon Fc-A treatment compared to the ethanol control treatments (**Fig 5A and 5B; Video S1**). GCaMP3 fluorescence between Fc-A- or ethanol-treated plants became distinct around 20 min after the treatments (**Fig 5C**), and the maximum changes of GCaMP3 fluorescence were significantly different between Fc-A- and ethanol-treated plants (**Fig 5D**), indicating that the PM H^+^-ATPase activation by Fc-A invoked the elevation of cytosolic Ca^2+^. These results imply that PM H^+^-ATPase promotes directly or indirectly the elevation of the cytosolic Ca^2+^ and induces the cell wall-related gene expression (**Fig 6**).

**Figure 5.**
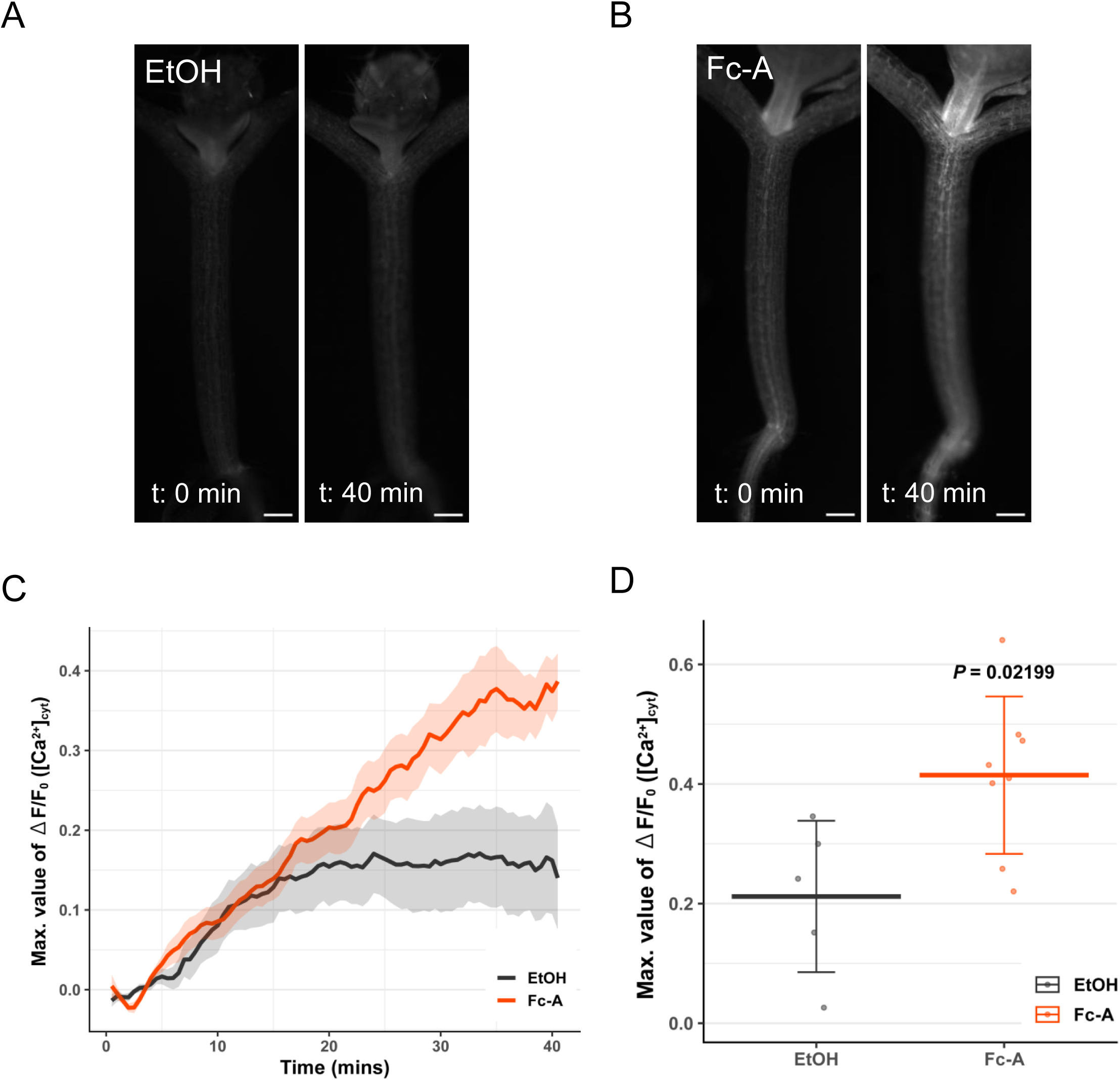
Cytosolic Ca^2+^ level changes upon Fc-A treatment. A) and B) shows a representative image of cytosolic Ca^2+^ monitoring in the hypocotyl of *p35S*::*GCaMP3*/ Col-0, before the treatment and 40 min after treatment. A) EtOH treated seedling and B) Fc-A treated seedling. Scale bars: 200 µm. C) The measured fluorescence of cytosolic Ca^2+^ biosensor normalised to the fluorescence before the treatment (ΔF/F_0_) along the time after the treatments; the line and the shade represent the mean ± S.E. in each conditions. *n* = 6−7. D) The comparison of maximum difference (Maximum ΔF/F_0_) in the measurement of C); Each point represents one biological replicates. Crossbars and error bars represent the mean ± S.D. in each conditions. The *P* value was determined by Welch’s *t* test.

**Figure 6.**
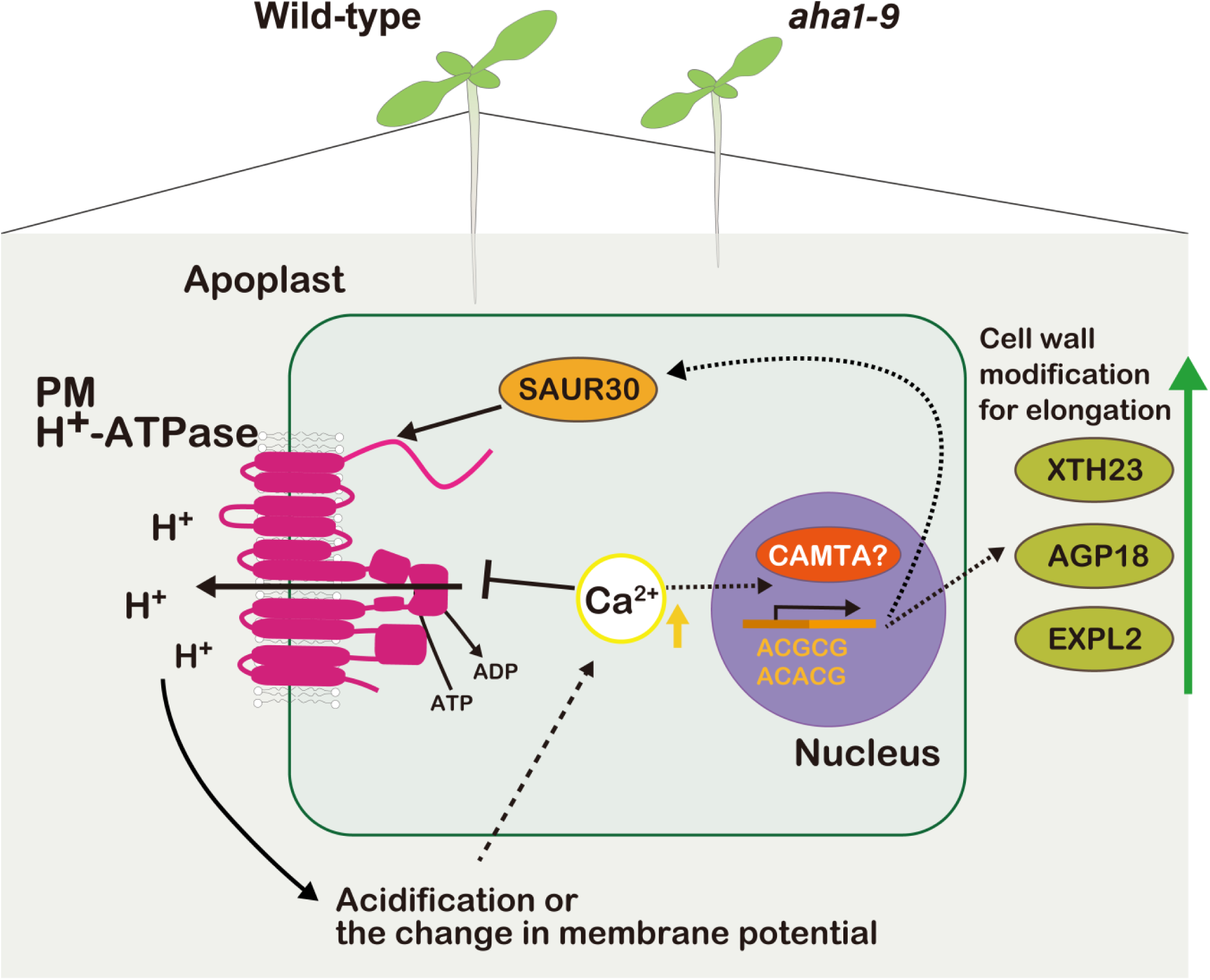
Hypothetical models of PM H^+^-ATPase activation triggered transcriptome change in photosynthetic tissues, i.e. green seedlings. When PM H^+^-ATPase is activated, the apoplast becomes acidic and the membrane potential across PM changes. These changes induce the cytosolic Ca^2+^ elevation with an unknown mechanism and the cytosolic Ca^2+^ elevation promotes the CAMTA transcription factor. CAMTA induces the expression of cell wall-related genes, resulting in the modification of cell wall properties that promote the elongation of cells. In addition, the elevation of Ca^2+^ reduces the PM H^+^-ATPase transiently and promotes the expression of an activator of PM H^+^-ATPase, SAUR30, which may fine-tune the activity of PM H^+^-ATPase.

## Discussion

### PM H^+^-ATPase promotes green seedlings’ growth by increasing the expression of cell wall modification-related genes

The idea of the acid growth has been accepted for a long time in plant science, while the model is mainly established in non-photosynthetic tissues, etiolated seedlings and roots. Here, we investigated how PM H^+^-ATPase activation promotes the growth of green seedling shoots. Our study showed that the growth of photosynthetically active green seedlings is also promoted by PM H^+^-ATPase, the primary source of “acid” in the apoplast. Our transcriptome analysis revealed that global transcript changes were induced upon PM H^+^-ATPase activation, and the elevation of cytosolic Ca^2+^ induces the changes in transcript levels. The induction of gene expression upon PM H^+^-ATPase activation was enriched in cell wall-related genes. Therefore, the induction of cell wall-related genes supports shoot growth by changing cell wall properties (**Fig 6**).

### PM H^+^-ATPase activation and elevation of cytosolic Ca^2+^

Elevation of cytosolic Ca^2+^ induces the Ca^2+^-dependent activation of CAMTA transcription factors (Finkler *et al*., 2007; Darwish *et al*., 2022). The Fc-A treatment induced the elevation of cytosolic Ca^2+^ in the hypocotyl of green seedlings (**Fig 5**). However, how the activation of PM H^+^-ATPase causes the elevation of cytosolic Ca^2+^ remains elusive. In other words, the connection between pH signalling and Ca^2+^ signalling has been a great target of plant science and discussed for decades (Li *et al*., 2021a). It is suggested that the increase of cytosolic pH, cytosolic alkalisation, invoke the elevation of Ca^2+^ in guard cells (Li *et al*., 2021a), as well as the apoplast acidification via exposure to low pH buffer also induces the Ca^2+^ signalling in root (Koyama *et al*., 2001). In contrast, some studies illustrate that the decrease of cytosolic pH upon wounding in leaves induces the elevation of Ca^2+^ (Behera *et al*., 2018), and a decrease of cytosolic pH via external application of ATP or glutamate induces the spike of Ca^2+^ in the root (Waadt *et al*., 2020). Our study applied the activator of PM H^+^-ATPase to green seedlings, managing constitutive pumping of the H^+^ from cytosol into apoplast, which leads to an increase of cytosolic pH and a decrease of apoplastic pH concurrently. The pH change across the PM by application of Fc-A is therefore, expected to be physiologically different from other previous studies using the external application of low pH buffer. In line with the hypothesis above, our RNA-seq analysis also differentiates the amplitude of response between Fc-A- and low pH treatment (**Fig 2**), suggesting that the activation of the PM H^+^-ATPase has a more substantial impact on global transcriptome than the application of other external stimuli. In addition, our Ca^2+^ monitoring assay showed the gradual elevation of cytosolic Ca^2+^ and became significantly different at 40 min after the treatment (**Fig 5**), while other studies with a state-of-art engineered genetically encoded biosensor mainly focus on the quick and steep Ca^2+^ elevation within 5 to 10 min after the treatments (Waadt *et al*., 2020; Li *et al*., 2021a). The difference of Ca^2+^ elevation pattern between this study and other studies may come from the tissue specificity and unique application of PM H^+^-ATPase activator. Thus, this study may provide the novel insight into the cytosolic Ca^2+^ signalling mediated by pH changes. However, interpreting the connection between pH change and cytosolic Ca^2+^ spike requires detailed observation, considering the time frame, treatment, and different Ca^2+^ monitoring biosensor system. Therefore, further investigations are needed to reveal how and what type of Ca^2+^ spike is induced by the PM H^+^-ATPase activation.

### Complex regulation of PM H^+^-ATPase activity

Elevation of Ca^2+^ represses the activity of PM H^+^-ATPase in guard cells (Kinoshita *et al*., 1995). Some plant Ca^2+^ sensors, SCaBP1 and SCaBP3, participate in repressing PM H^+^-ATPase activity in seedlings via recruiting the PKS5 kinase (Fuglsang *et al*., 2007; Yang *et al*., 2019). Application of rapid alkalisation factor, RALFs, induces the Ca^2+^ spike and repression of PM H^+^-ATPase, leading to a quick and transient alkalisation of the apoplast in roots, possibly via FERRONIA-mediated posttranslational regulation of the PM H^+^-ATPase (Gjetting *et al*., 2020). The study of RALF also suggests that apoplast acidification induces the expression of RALFs, thus forming negative feedback regulation of PM H^+^-ATPase. In line with the RALF study, our RNA-seq confirmed that Fc-A treatment induced a putative *RALF*, *RALF-like 33* (AT4G15800; **Data S1**), as well as identified *SAUR30*, the positive feedback regulators (**Fig S5**), which has some CAMTA binding sites in the promoter (**Fig S5B**) and phylogenetically belongs to the same clade as Ca^2+^ responsive SAURs (Zhang *et al*., 2021). Taken together, we propose that, upon PM H^+^-ATPase activation, cytosolic Ca^2+^ elevation induces the quick repression of PM H^+^-ATPase activity by posttranslational modification and then finely modulates the proper activity of PM H^+^-ATPase via transcriptional induction of the expression of PM H^+^-ATPase positive and negative regulators.

### Novel perspective of acid growth model

In the acid growth model, the phytohormones, auxin and brassinosteroids, are the main activators of PM H^+^-ATPase in the hypocotyl of etiolated seedlings or roots (Wang *et al*., 2014; Miao *et al*., 2022), therefore, the acid growth is generally perceived as phytohormone-dependent phenomena. A recent study illustrates that the activation of PM H^+^-ATPase in photosynthetic tissues such as leaves is regulated by photosynthetic products (Okumura *et al*., 2016; Kinoshita *et al*., 2023). The hypocotyl elongation rate in photosynthetic seedlings is inhibited compared to etiolated seedlings due to the low phytohormone content and reduced sensitivity (Vert *et al*., 2008; Kurepin *et al*., 2011). However, the photosynthetic tissue growth, such as leaf expansion, is still maintained at a certain level. Therefore, it is possible that, in photosynthetic tissues, the acid growth model is supported by photosynthetic product-dependent activation of PM H^+^-ATPase as the light illumination as well as sucrose feeding to leaves indeed upregulated essential genes for cell wall modification but not auxin- or brassinosteroids-related genes (**Fig S3**). Together with this study’s results, we propose adding a new perspective on the acid growth model, shedding light on the impact of PM H^+^-ATPase activation in photosynthetic tissues, probably mediated by photosynthesis.

## Acknowledgements

We would like to thank Akiko Akama and Mikako Yamaguchi from the Center for Gene Research, Nagoya University for the support with next-generation sequencing (NextSeq550), and Mayuko Naganawa and Riko Hasegawa for experimental support. We appreciate a financial support and grants from the Ministry of Education, Culture, Sports, Science, and Technology, Japan (grant nos. 20H05687 and 20H05910 to T.K.) and grant-in-aid for Japan Society for the Promotion of Science Research Fellow (grant no. 19J20450 to S.N.K), and MEXT (WISE Program) “Graduate Program of Transformative Chem-Bio Research” in Nagoya University.

## Author contributions

S.N.K. and T.K. initiated and conceptualised the project. S.N.K. and K.Taki conducted most of the experiments except for the cDNA library preparation and RNA-seq. F.O. set up the calcium imaging system. J.O. purified the Fc-A. M.N. and Y.T. conducted RNA-seq and *in silico* promoter analysis. S.N.K. and Y.H. analysed and mapping the raw data from RNA-seq. Y.H. and K.Takahashi generated *gAHA1*/*aha1-9*. S.N.K. prepared the manuscript and figures. I.F. provided supervision and discussion. All authors reviewed the manuscript.

## Data availability

RNA-seq raw read data has been deposited in DDBJ Sequence Read Archive (DRA) with the accession number, BioProject: PRJDB18676.

---

**Figure S1.**
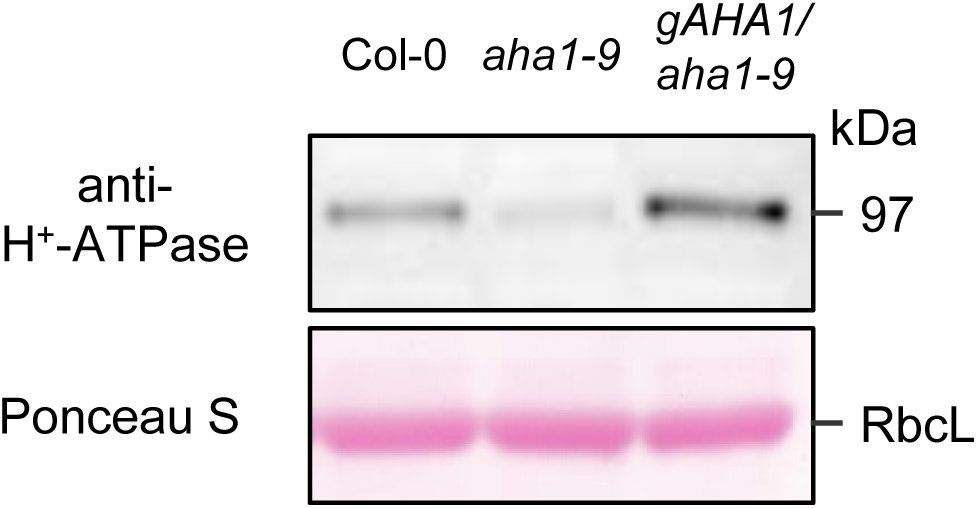
PM H^+^-ATPase abundance in the *aha1-9* mutant and complementation line, *gAHA1*/ *aha1-9*. A representative image of the Western blot analysis, confirming the complementation of PM H^+^-ATPase in *gAHA1*/ *aha1-9*. PM H^+^-ATPase abundance was detected by the PM H^+^-ATPase specific antibody and total loaded protein amount was visualised by the PonceauS staining of membrane.

**Figure S2.**
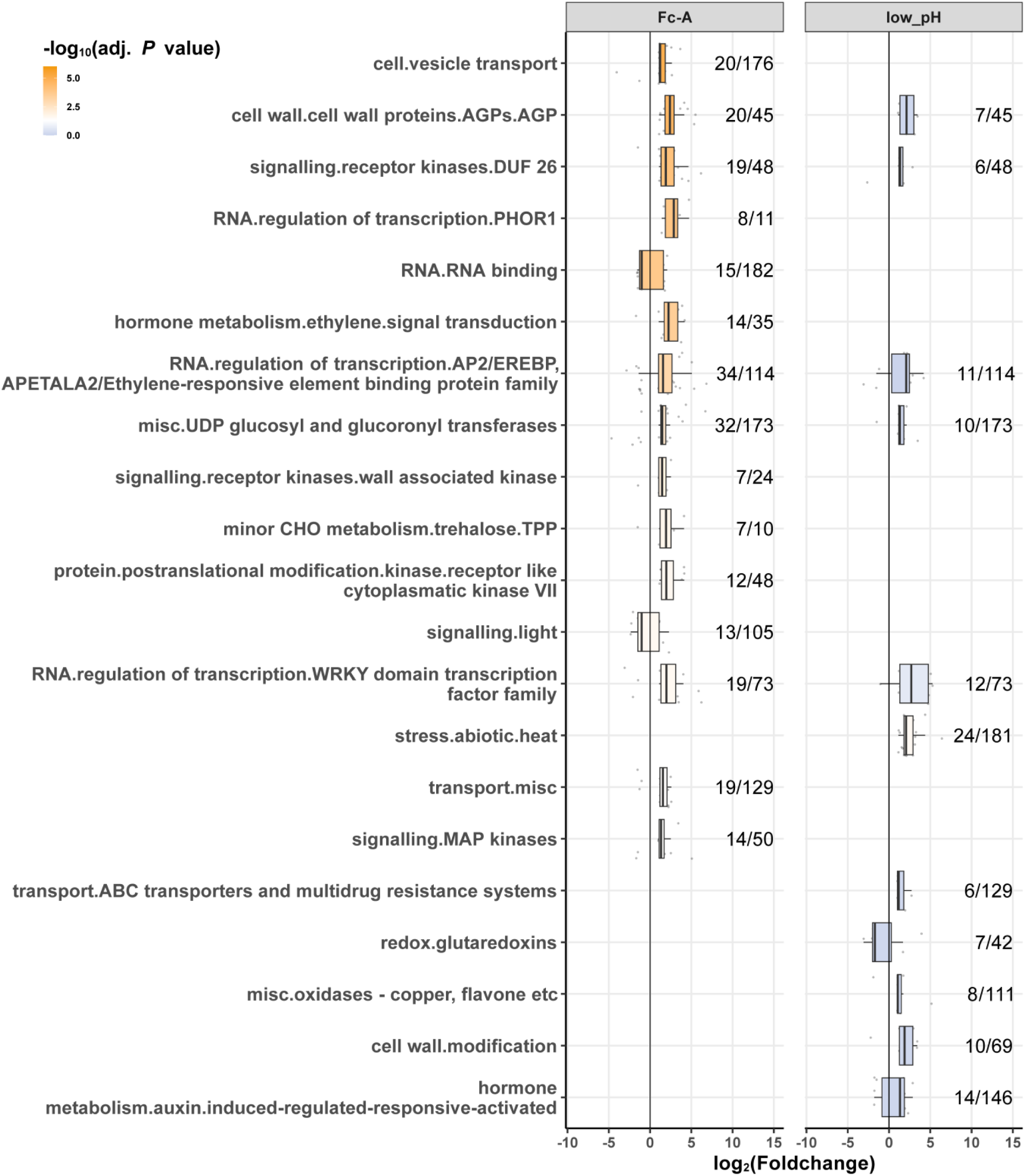
Foldchange of transcripts in specified MapMan groups. The boxplots represent the distribution of DEGs expression changes in functional category groups compared to the ethanol treated samples. The numbers beside the boxplots indicate the number of DEGs over the number of all registered genes in the category groups. Adjusted *P* values were determined by Mann-Whitney U test in MapMan.

**Figure S3.**
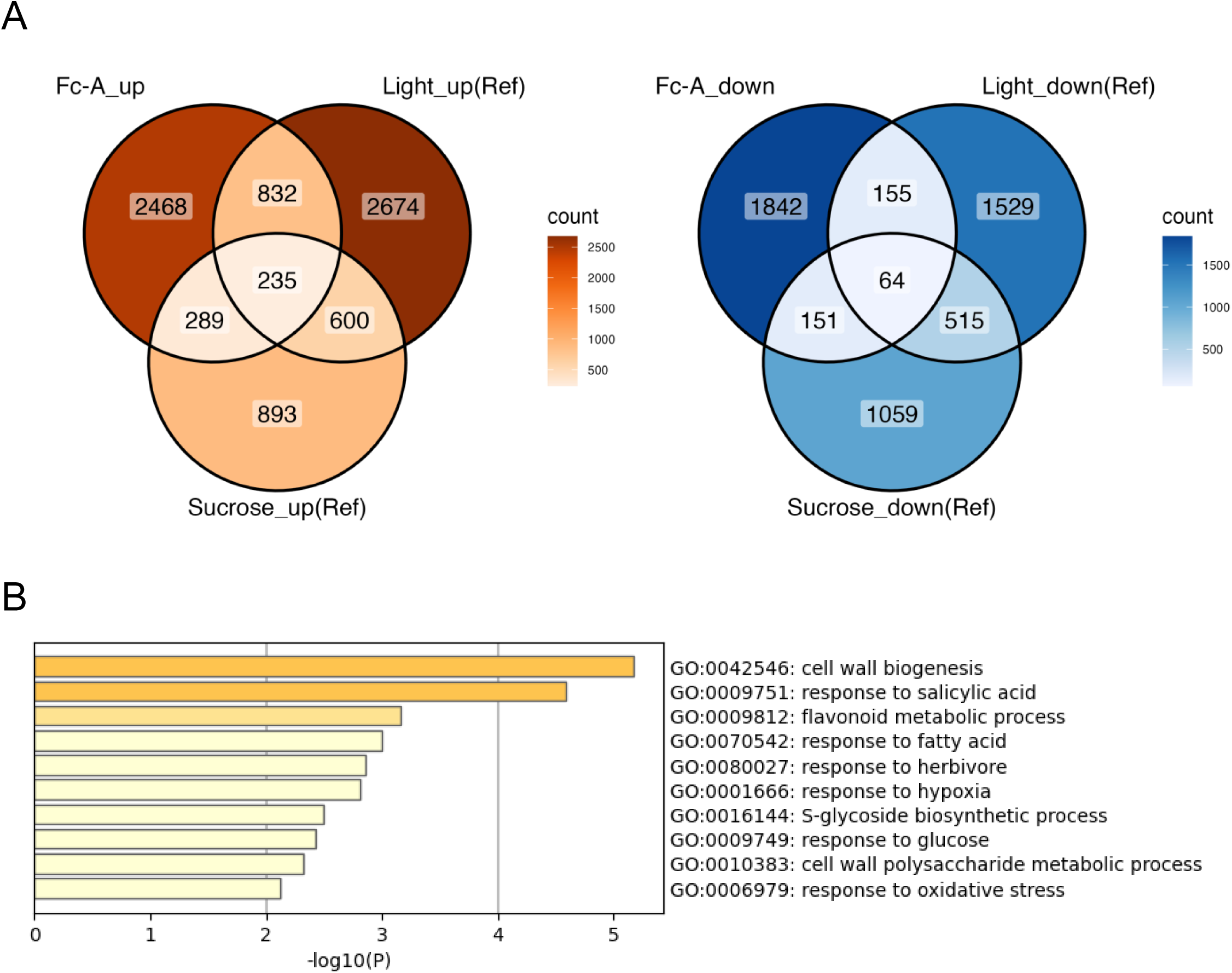
Comparison with photosynthate-dependent transcriptome change in leaves. A) Venn diagrams represent the numbers of upregulated transcripts (left; foldchange > 2) or downregulated transcripts (right; foldchange < −2) in Fc-A treated seedlings compared to ethanol treated seedlings, light illuminated leaves or sucrose supplemented leaves from Kinoshita et al. 2023. B) GO term enrichments of commonly upregulated transcripts (235 genes in the panel A).

**Figure S4.**
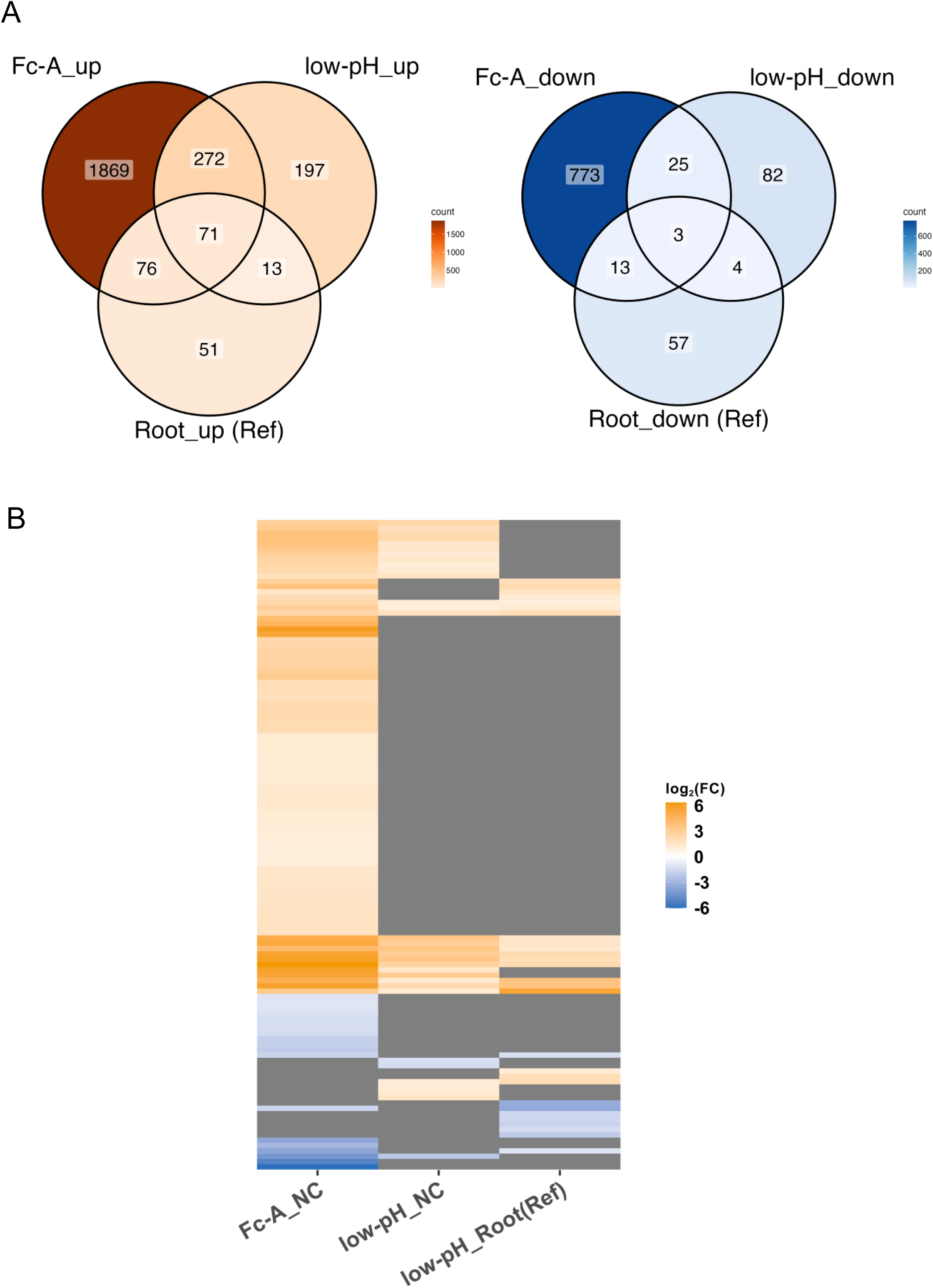
Comparison with low-pH treatment-dependent transcriptome change in roots. A) Venn diagram represent the numbers of differentially expressed genes (DEGs; |foldchange| > 2, FDR < 0.05) in Fc-A treated or low pH media treated seedlings compared to ethanol treated seedlings and Ref. data obtained from microarray results from Lager et al. 2010 using low-pH treated roots of Arabidopsis. On the left are upregulated DEGs and on the right are downregulated DEGs. B) Heatmap presents the foldchange (FC) of cell wall-related genes in this study and Ref. data from Lager et al. 2010. Negative control (NC) is EtOH treated seedling shoots. Grey colour in heatmap represent either not significant or not detected in the datasets.

**Figure S5.**
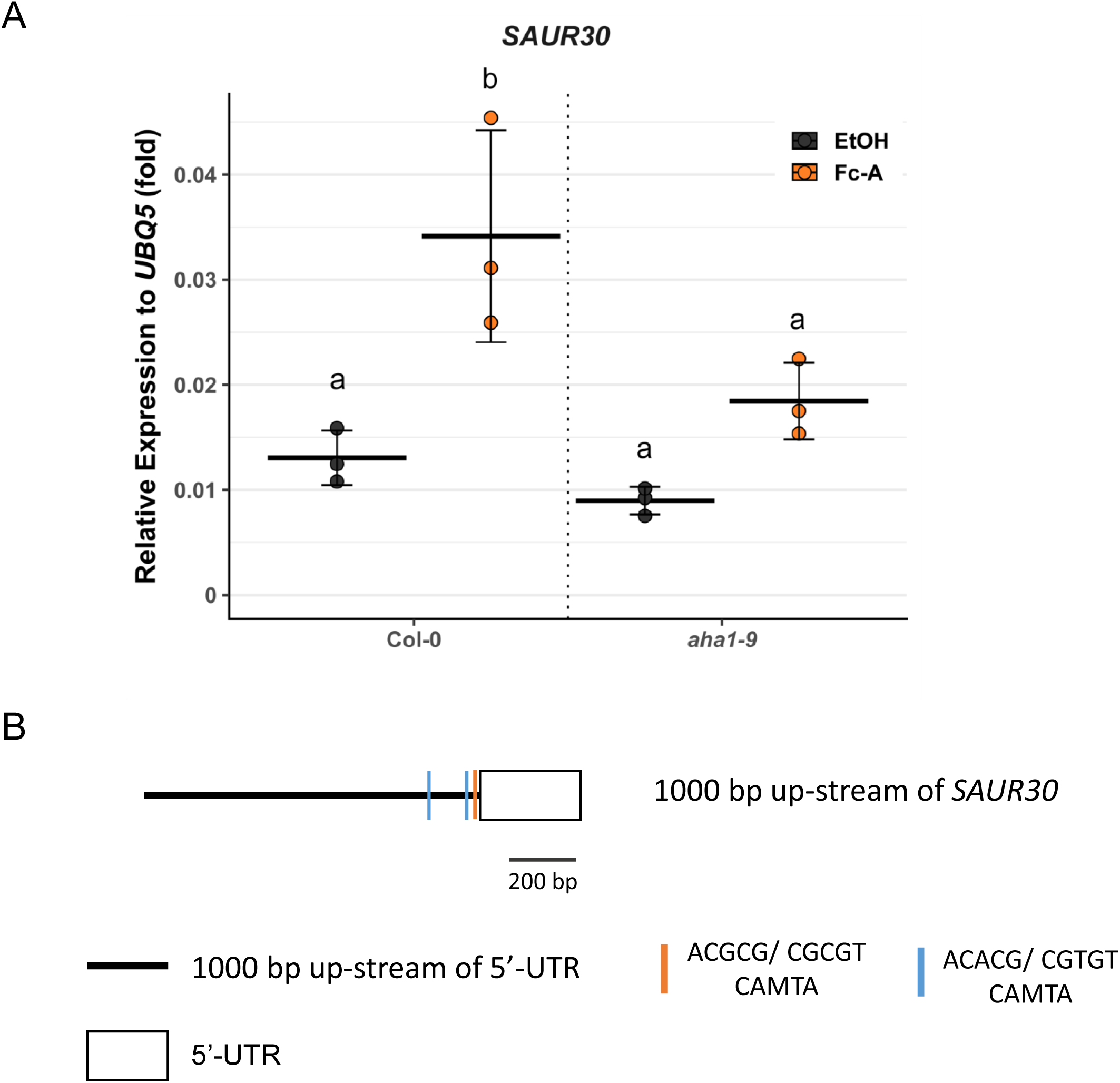
*SAUR30* expression change in wildtype and *aha1-9*. A) The relative expression of *small auxin up RNA 30* (*SAUR30*) to ubiquitous *UBQ5* in wildtype (Col-0) and *aha1-9,* determined by RT-qPCR. Each point represents one biological replicate. Crossbars and error bars represent the mean ± S.D. in each condition. Different letters above error bars indicate the significant difference in gene expression, determined by one-way ANOVA with Tukey HSD. B) Schematic diagram of the pentamer positions in the promoter region of *SAUR30* genes. Black line represents the 1000 bp upstream of 5’-UTR; blue narrow line and orange narrow line indicate the position of GCGC box, ACGCG and ACACG pentamer, respectively; white box represents the position of 5’-UTR.

**Figure S6.**
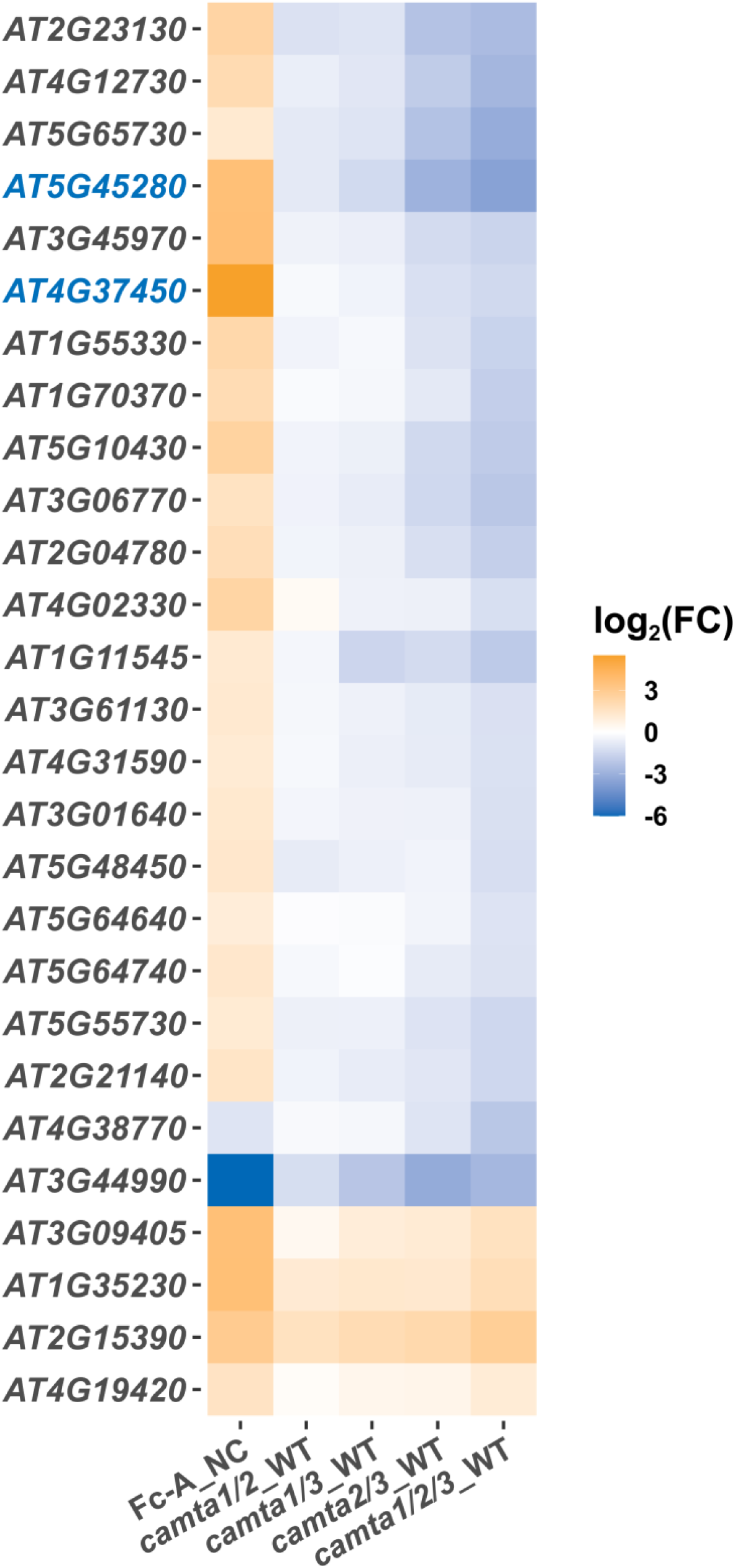
CAMTA-dependent expression profile of cell wall-related genes. Heatmap presents the foldchange (FC) of cell wall-related genes in this study and Ref. data, *camta* mutant analysis from Kim et al. 2013. Negative control (NC) is EtOH treated seedling shoots. Only detected cell wall-related genes both in this study and Kim et al. 2013 are listed. Blue colour of the AGI number indicates the genes tested in RT-qPCR of this study.

**Figure S7.**
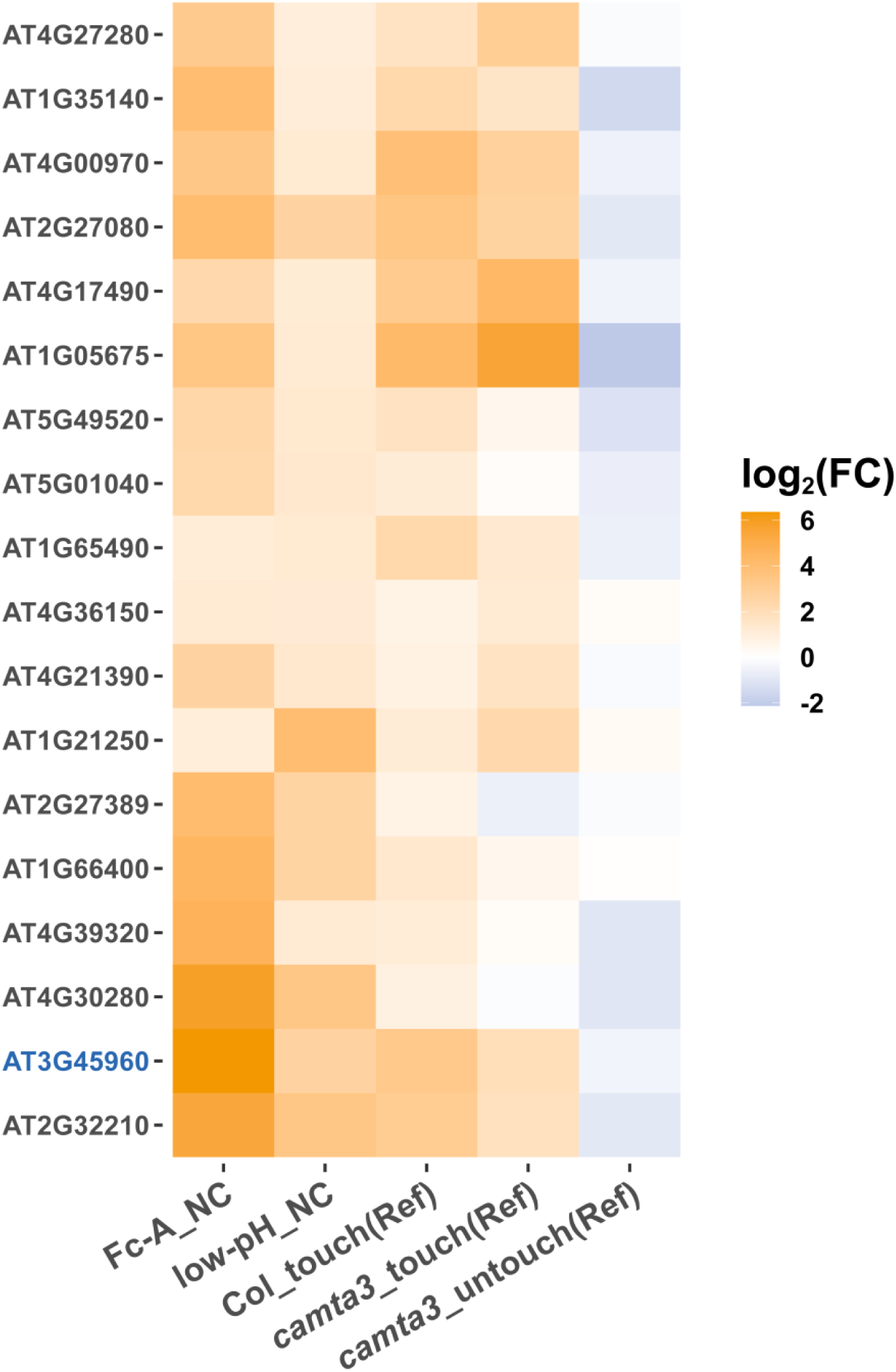
*CAMTA3*-dependent touch-induced expression patterns. Heatmap presents the foldchange (FC) of touch-induced genes in this study and Ref. data, using touch treatment and *camta* mutant from Darwish et al. 2022. Negative control (NC) is EtOH treated seedling shoots. The foldchange of gene expression in Darwish et al 2022 are comparison to the untouched Col-0 sample. Only detected genes both in this study and in Darwish et al 2022 are listed. Blue colour of the AGI number indicates the genes tested in RT-qPCR of this study.

